# Quality Matters: Biocuration Experts on the Impact of Duplication and Other Data Quality Issues in Biological Databases

**DOI:** 10.1101/788034

**Authors:** Qingyu Chen, Ramona Britto, Ivan Erill, Constance J. Jeffery, Arthur Liberzon, Michele Magrane, Jun-ichi Onami, Marc Robinson-Rechavi, Jana Sponarova, Justin Zobel, Karin Verspoor

## Abstract

The volume of biological database records is growing rapidly, populated by complex records drawn from heterogeneous sources. A specific challenge is duplication, that is, the presence of redundancy (records with high similarity) or inconsistency (dissimilar records that correspond to the same entity). The characteristics (which records are duplicates), impact (why duplicates are significant), and solutions (how to address duplication), are not well understood. Studies on the topic are neither recent nor comprehensive. In addition, other data quality issues, such as inconsistencies and inaccuracies, are also of concern in the context of biological databases. A primary focus of this paper is to present and consolidate the opinions of over 20 experts and practitioners on the topic of duplication in biological sequence databases. The results reveal that survey participants believe that duplicate records are diverse; that the negative impacts of duplicates are severe, while positive impacts depend on correct identification of duplicates; and that duplicate detection methods need to be more precise, scalable, and robust. A secondary focus is to consider other quality issues. We observe that biocuration is the key mechanism used to ensure the quality of this data, and explore the issues through a case study of curation in UniProtKB/Swiss-Prot as well as an interview with an experienced biocurator. While biocuration is a vital solution for handling of data quality issues, a broader community effort is needed to provide adequate support for thorough biocuration in the face of widespread quality concerns.

## Introduction

The major biological databases represent an extraordinary collective volume of work. Diligently built up over decades and comprised of many millions of contributions from the biomedical research community, biological databases provide worldwide access to a massive number of records (also known as *entries*) [1]. Starting from individual laboratories, genomes are sequenced, assembled, annotated, and ultimately submitted to primary nucleotide databases such as GenBank [2], ENA [3], and DDBJ [4] (collectively known as INSDC). Translations of those nucleotide records, protein records, are deposited into central protein databases such as the UniProt KnowledgeBase (UniProtKB) [5] and the Protein Data Bank [6]. Sequence records are further accumulated into different databases for more specialised purposes: RFam [7] and PFam [8] for RNA and protein families respectively, such as DictyBase [9] and PomBase [10] for model organisms, ArrayExpress [11] and GEO [12] for gene expression profiles. These databases are selected as examples; the list is not intended to be exhaustive. However, they are representative of biological databases that have been named in the “golden set” of the 24th Nucleic Acids Research database issue. The introduction of that issue highlights the databases that “consistently served as authoritative, comprehensive, and convenient data resources widely used by the entire community and offer some lessons on what makes a successful database” [13]. The associated information about sequences is also propagated into non-sequence databases, such as PubMed (https://www.ncbi.nlm.nih.gov/pubmed/) for the scientific literature, or GO [14] for function annotations. Those databases in turn benefit individual studies, many of which use these public available records as the basis for their own research.

Inevitably, given the scale of these databases, some submitted records are redundant [15], inconsistent [16], inaccurate [17], incomplete [18], or outdated [19]. Such quality issues can be addressed by manual curation, with the support of automatic tools, and by processes such as reporting of the issues by contributors detecting mistakes. Biocuration plays a vital role in biological database curation [20]. It de-duplicates database records [21], resolves inconsistencies [22], fixes errors [17], and resolves incomplete and outdated annotations [23]. Such curated records are typically of high quality and represent the latest scientific and medical knowledge. However, the volume of data prohibits exhaustive curation, and some records with those quality issues remain undetected.

In other work, we (Chen, Verspoor, and Zobel) have explored a particular form of quality issue, which we have characterized as *duplication* [24,25]. As described in that work, duplicates are characterized in different ways in different contexts, but they can be broadly categorized as *redundancies* or *inconsistencies*. The perception of a pair of records as duplicates depends on the task. As we wrote in previous work, *“a pragmatic definition for duplication is that a pair of records A and B are duplicates if the presence of A means that B is not required, that is, B is redundant in the context of a specific task or is superseded by A.”* [24]. Many such duplicates have been found through curation, but the prevalence of undetected duplicates is unknown, as is the accuracy and sensitivity of automated tools for duplicate or redundancy detection. Other work has explored the detection of duplicates, but often under assumptions that limit the impact. For example, some researchers have assumed that similarity of genetic sequence is the sole indicator of redundancy, whereas in practice some highly similar sequences may represent distinct information and some rather different sequences may in fact represent duplicates [26]. We detail the notion and impacts of duplication in the next section.

### Authors’ contributions

In this work, a main focus is to explore the characteristics, impacts, and solutions to duplication in biological databases; a secondary focus is to further investigate other quality issues. We present and consolidate the opinions of over 20 experts and practitioners on the topic of duplication and other data quality issues, via a questionnaire-based survey. To address different quality issues, we introduce biocuration as a key mechanism for ensuring the quality of biological databases. To our knowledge, there is no one-size-fits-all solution even to a single quality issue [27]. We thus explain the complete UniProtKB/Swiss-Prot curation process, via a descriptive report and an interview with its curation team leader, which provides a reference solution to different quality issues. Overall, the observations on duplication and other data quality issues highlight the significance of biocuration in data resources, but a broader community effort is needed to provide adequate support to facilitate thorough biocuration.

### The notion and impact of duplication

Our focus is on database records – that is, entries in structured databases – not on biological processes such as gene duplication. Superficially, the question of what constitutes an *exact duplicate* in this context can seem obvious: two records that are exactly identical in both data (*e.g*., sequence) and annotation (*e.g*., meta-data including species and strain of origin) are duplicates. However, the notion of duplication varies. We demonstrate a generic biological data analysis pipeline involving biological databases and illustrate different notions of duplication.

**Figure 1** shows the pipeline; we explain the three stages of the pipeline using the databases managed by the UniProt Consortium (http://www.uniprot.org/) as examples.

**Figure 1.**
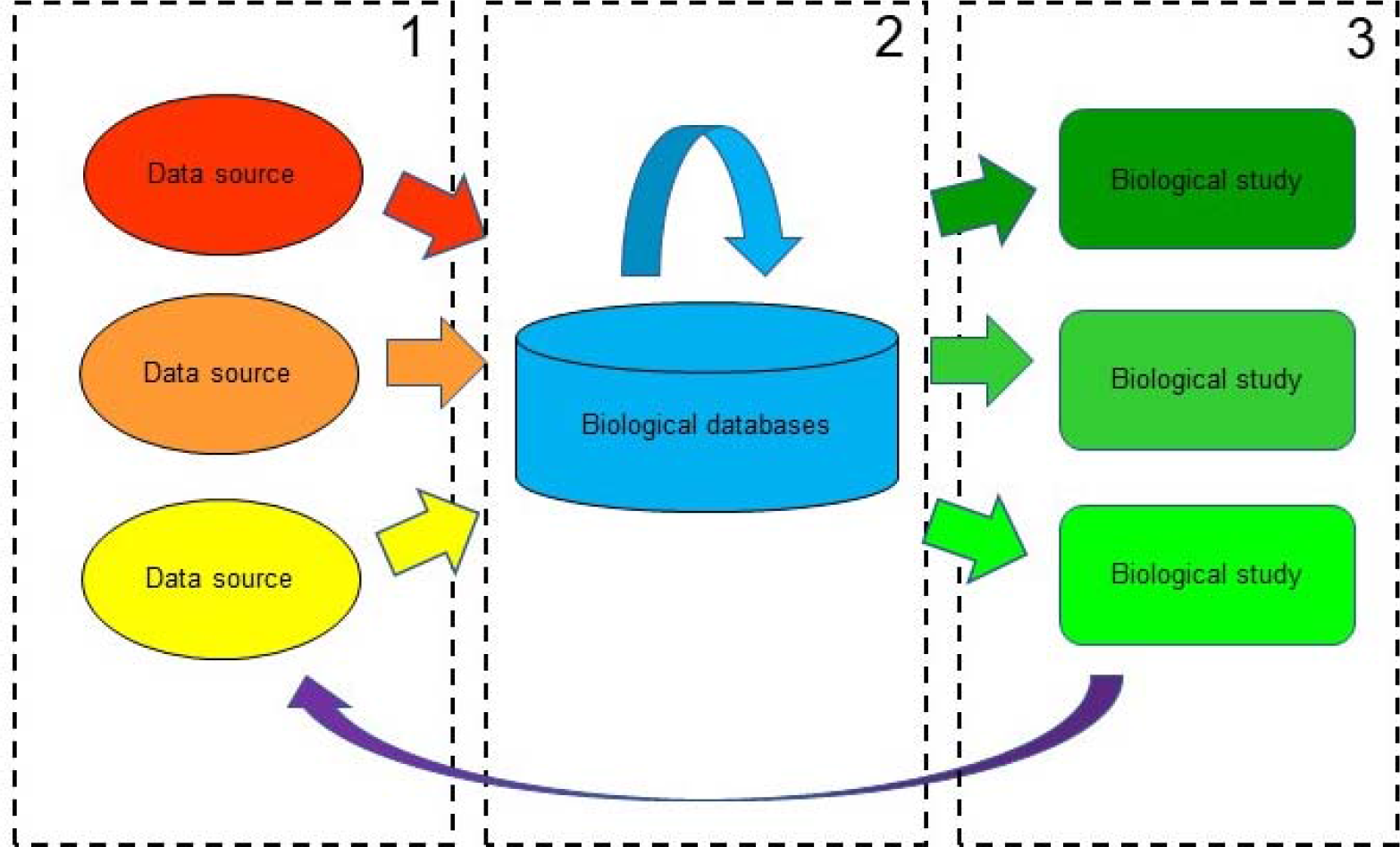
Biological analysis pipeline. Three stages of a biological analysis pipeline, heavily involving biological databases, are presented.

**“pre-database” stage**: records from various sources are submitted to databases. For instance, UniProt protein records come from translations of primary INSDC nucleotide records (directly submitted by researchers), direct protein sequencing, gene prediction and other sources (http://www.uniprot.org/help/sequence_origin).

**“within database” stage**: database curation, search, and visualisation. Records are annotated in this stage, automatically (UniProtKB/TrEMBL) or through curation (UniProtKB/Swiss-Prot). Biocuration plays a vital role at this stage. For instance, UniProt manual curation not only merges records and documents discrepancies, it also annotates the records with biological knowledge drawn from the literature [28]. Also, the databases need to manage the records for search and visualisation purposes [29]. During this stage, UniProt undertakes extensive cross-referencing by linking hundreds of databases to provide centralized knowledge and resolve ambiguities [30].

**“post-database” stage**: record download, analysis, and inference. Records are downloaded and analysed for different purposes. For instance, both UniProtKB records and services have been extensively used in the research areas of biochemistry and molecular biology, biotechnology and computational biology, according to citation patterns [31]. The findings of studies may in turn contribute to new sources.

Duplication occurs in all of these stages, but its relevance varies. Continuing with the UniProt example, the first stage primarily concerns *entity duplicates* (often referred to as *true duplicates*): records that correspond to the same biological entities regardless of whether there are differences in the content of the database records. Merging those records into a single entry is the first step in UniProtKB/Swiss-Prot manual curation [28]. The second stage primarily concerns *near-identical duplicates (*often referred to as *redundant records)*: the records may not refer to the same entities, but nevertheless have high similarity. UniProt has found those records lead to uninformative BLAST search results (http://www.uniprot.org/help/proteome_redundancy). The third stage primarily concerns *study-dependent duplicates*: studies may further de-duplicate sets of records for their own purposes. For instance, studies on secondary protein structure prediction may further remove protein sequences at a 75% sequence similarity threshold [32]. This clearly shows that the notion of duplication varies and in general has two characteristics: *redundancy and inconsistency*. Thus it is critical to understand their characteristics, impacts, and solutions.

We have found numerous discussions of duplicates in the previous literature. As early as in 1996, Korning et al. [33] observed duplicates from the GenBank *Arabidopsis thaliana* dataset when curating those records. The duplicates were of two main types: the same genes that were submitted twice (either by the same or different submitters), and different genes from the same gene family that were similar enough that only one was retained. Similar cases were also reported by different groups [21, 34–36]. Recently, the most significant case was the duplication in UniProtKB/TrEMBL [15]: in 2016, UniProt removed 46.9 million records corresponding to duplicate proteomes (for example, over 5.9 million of these records belong to 1,692 strains of *Mycobacterium tuberculosis*). They identified duplicate proteome records based on three criteria: belonging to the same organisms; sequence identity of over 90%; and the proteome ranks designed by biocurators (such as whether they are Reference proteome and the annotation level).

As this history shows, investigation of duplication has persisted for at least 20 years. Considering the type of duplicates, as the above discussion illustrates, duplication appears to be richer and more diverse than was originally described (we again note the definition of ‘duplication’ we are following in this paper, which includes the concept of redundancy). This motivates continued investigation of duplication.

An underlying question is: does duplication have positive or negative impact? There has been relatively little investigation of the impact of duplication, but there are some observations in the literature: (1) “The problem of duplicates is also existent in genome data, but duplicates are less interfering than in other application domains. Duplicates are often accepted and used for validation of data correctness. In conclusion, existing data cleansing techniques do not and cannot consider the intricacies and semantics of genome data, or they address the wrong problem, namely duplicate elimination.” [38]; (2) “Biological data duplicates provide hints of the redundancy in biological datasets … but rigorous elimination of data may result in loss of critical information.” [34]; (3) “The bioinformatics data is characterized by enormous diversity matched by high redundancy, across both individual and multiple databases. Enabling interoperability of the data from different sources requires resolution of data disparity and transformation in the common form (data integration), and the removal of redundant data, errors, and discrepancies (data cleaning).” [39]. Thus the answers to questions on the impact of duplicates are not clear. The above views are inconsistent, are opinions rather than conclusions drawn from studies, and are not supported by extensive examples. Moreover, they are not recent, and may not represent the current environment. Answering the question of the impact of duplications requires a more comprehensive and rigorous investigation.

### From duplication to other data quality issues

Biological sources suffer from data quality issues other than duplication. We summarise diverse biological data quality issues reported in the literature: inconsistencies (such as conflicting results reported in the literature) [22], inaccuracies (such as erroneous sequence records and wrong gene annotations) [40–42], incompleteness (such as missing exons and incomplete annotations) [38, 40] and outdatedness (such as out-dated sequence records and annotations) [41]. This shows that while duplication is a primary data quality issue, other quality issues are also of concern. Collectively, there are five primary data quality issues: duplication, inconsistency, inaccuracy, incompleteness and outdatedness identified in general domains [43]. It is thus also critical to understand what quality issues have been observed and how they impact database stakeholders under the context of biological databases.

### Practitioner viewpoint: survey questions

Studies on data quality broadly take one of three approaches: domain expertise, theoretical or empirical. The first is opinion-based: accumulating views from (typically a small group of) domain experts [44–46]. For example, one book summarises opinions from domain experts on elements of spatial data quality [44]. The second is theory-based: inference of potential data quality issues from a generic process of data generation, submission, and usage [47–49]. For example, a data quality framework was developed by inferring the data flow of a system (such as input and output for each process) and estimating the possible related quality issues [47]. The third is empirically based: analysis of data quality issues in a quantitative manner [50–52]. For example, an empirical investigation on what data quality means to stakeholders was performed via a questionnaire [50]. Each approach has its own strengths and weaknesses; for example, opinion-based studies represent high domain expertise, but may be narrow due to the small group size. Quantitative surveys in contrast have a larger number of participants, but the level of expertise may be relatively lower.

Our approach integrates opinion-based and empirical-based approaches: the study presents opinions from domain experts; but the data was gathered via a questionnaire; the survey questions are provided in the Supplementary Material File S1. We surveyed 23 practitioners on the questions of duplicates and other general data quality issues. These practitioners are from diverse backgrounds (including experimental biology, bioinformatics, and computer science), with a range of affiliation types (such as service providers, universities, or research institutes) but all have domain expertise. These practitioners include senior database staff, project and lab leaders, and biocurators. The publications of the participants are directly relevant to databases, data quality and curation; as illustrated by some instances [10, 15, 28, 53–69]. They were selected by personal approach at conferences and in a small number of cases by email; most of the practitioners were not known to the originating authors (Chen, Verspoor, Zobel) before this study.

A limitation is that the small participant size may mean that we have collected unrepresentative opinions. However, the community of biocuration is small and the experience represented by these 23 is highly relevant. A 2012 survey conducted by the International Society of Biocuration (ISB) had 257 participants [67]. Of those 257 participants, 57% of them were employed in short-term contracts and only 9% were principal investigators. A similar study initiated by the BioCreative team had only 30 participants, including all the attendees of the BioCreative conference in that year [68]. Therefore, the number of participants of this study reflects the size of the biocuration community; moreover, the relatively high expertise ensures the validity of the opinions.

The survey asked three primary questions about duplication: (1) *What* are duplicates? We asked practitioners what records they think should be regarded as duplicated; (2) *Why* care about duplicates? We asked practitioners what impact duplicates have; (3) *How* to manage duplicates? We asked practitioners whether, and how, duplicates should be resolved.

In detail, the questions and their possible responses were as follows:

#### Defining duplicate records (The ‘what’ question)

We provided five options for experts to select: (1) Exact duplicate records: two or more records are exactly identical; (2) Near identical duplicates: two or more records are not identical but similar; (3) Partial or fragmentary records: one record is a fragment of another; (4) Duplicate records with low similarity: records have relatively low similarity but belong to the same entity; (5) Other types: if practitioners also consider other cases as duplicates.

Respondents were asked to comment on their choices. We also requested examples to support the choice of options 4 or 5, given that in our review of the literature we observed that the first three options were prevalent [70, 71]. Option 1 refers to exact duplicates, option 2 refers to (highly) similar or redundant records or to some quantitative extent, records share X% similarity, option 3 refers to partial or incomplete records, and option 4 refers to entity duplicates that are inconsistent. The “Other types” option provides capture of remaining types of duplicates.

#### Quantifying the impacts of duplication (The ‘Why’ question)

We asked in two steps: first, whether respondents believed that duplicates have impact. The second question was presented only if the answer to the first was yes. It is used to comment on positive and negative impacts respectively. We also asked respondents to explain their opinion or give examples.

#### Addressing duplication (The ‘How’ question)

We offered three subquestions: (1) Do you believe that duplicate detection is useful/needed? (2) Do you believe that current duplicate detection methods/software are sufficient to satisfy your requirements? (We also asked respondents to explain what they expected if they selected ‘no’.) (3) How would you prefer that duplicate records be handled? These were the suggested options: label and remove duplicates, label and make duplicates obsolete, label but leave duplicates active, and other solutions.

### Practitioner viewpoints: summary

In this section, we present the survey results on duplication and other quality issues.

#### Duplication: practitioners’ opinions

The responses are summarized below, in the same order as the three primary questions. For each question, we detail the response statistics, summarise the common patterns, augmented by detailed responses, and draw conclusions.

The views on *what are duplicates* are summarised in **Figure 2**. Out of 23 practitioners, 21 have made a choice by selecting at least one option. While the other two did not select any options, they have considered that duplicates have impacts for later questions. We therefore do not regard the empty responses as an opinion that duplication does not exist; rather simply do not track the response in this case.

**Figure 2.**
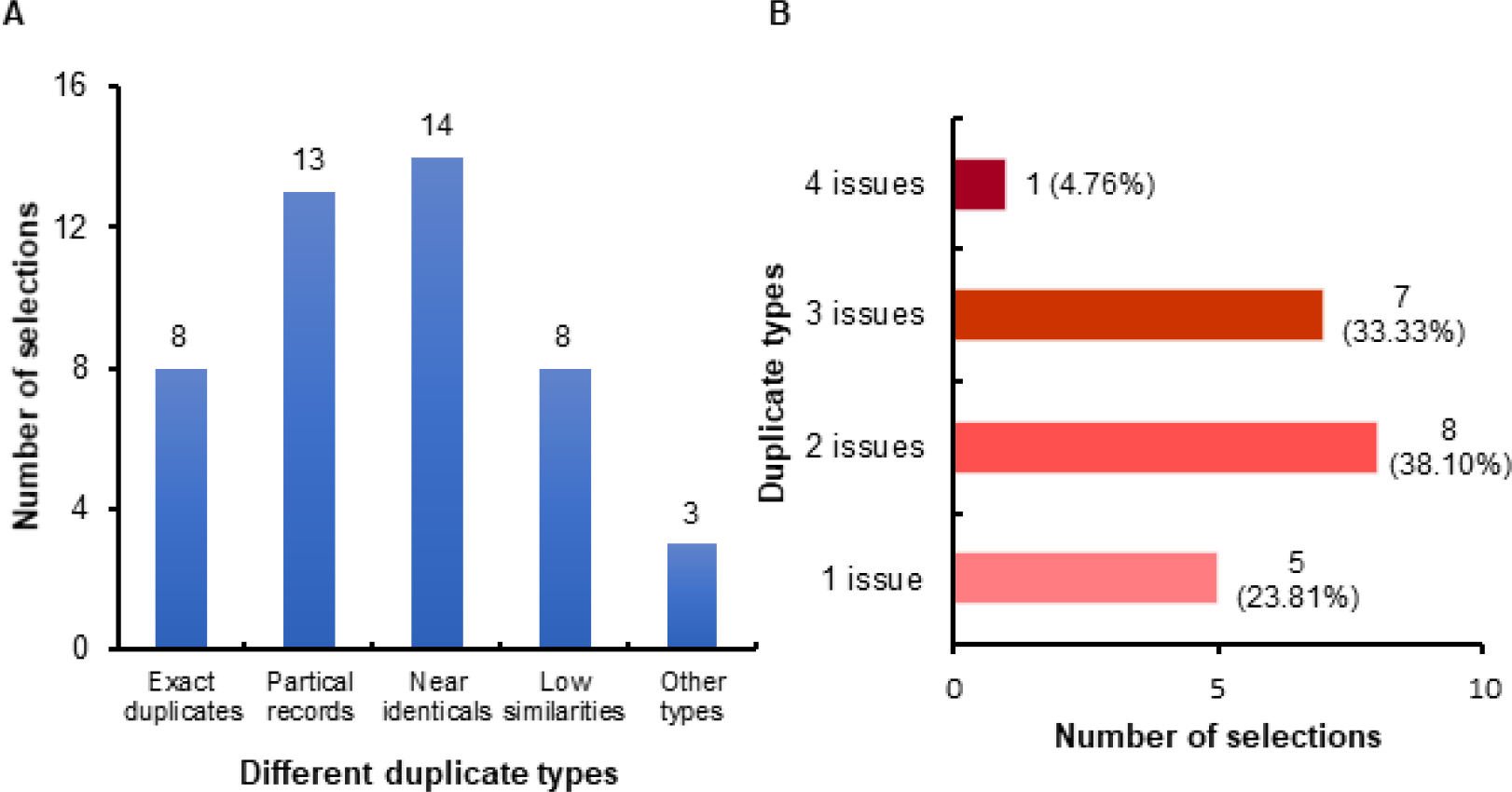
Characteristics of duplicate records. What are duplicates? The X-axis shows different duplicate types; the Y-axis shows the associated number of participants who selected that type.

The results show (1) all types of duplicates have been observed by some of practitioners, but none is universal. The commonest type is *similar record*, which was selected by over half of the respondents; but the other types (*exact duplicates, partial records*, and *low similarity duplicates*) were also selected by at least a third of the respondents. Three of them considered *other duplicate* types, and (2) more than 80% of respondents indicated that they have observed at least two types.

Also recall that existing literature rarely covers the fourth type of duplication – that is, relatively different records that should in fact be considered as duplicates. However, close to 40% of respondents acknowledge having seen such cases and further point out that identifying them requires significant manual effort. The following summarises several cases (each identified by respondent ID, tabulated at the end of this paper).

*Low similarity duplicates within a single database*. Representative comments are “We have such records in ClinVar [64]. We receive independent submissions from groups that define variants with great precision, and groups that define the same variant in the same paper, but describe it imprecisely. Curators have to review the content to determine identity.” [R19] and “Genomes or proteomes of the same species can often be different enough even they are redundant.” [R24]

##### Low similarity duplicates in databases having cross-references

Representative comments are “Protein-Protein Interaction databases: the same publication may be in BioGRID [72] annotated at the gene level and in one of the IMEx databases (http://www.imexconsortium.org/) annotated at the protein level.” [R20] and “Also secondary databases import data (e.g. STRING sticking to the PPI example) but will only import a part of what is available.” [R20].

##### Low similarity duplicates in databases having the same kinds of contents

For instance, “Pathway databases (KEGG^29^, Reactome^30^, EcoCyc^31^ etc) tend to look at same pathways but are open to curator interpretation and may differ.” [R20]

The results of the “why care about duplicates” question are shown in **Figure 3**. All practitioners made a choice. Most (21 out of 23) believe that duplication does matter. Moreover, 19 out of 21 experts weighted on potential impact of duplicates: only one believed that the impact is purely positive, compared to 8 viewing it solely negative; the remaining 10 thought the impact has both positive and negative sides. We assembled all responses on impacts of duplicates as follows below.

**Figure 3.**
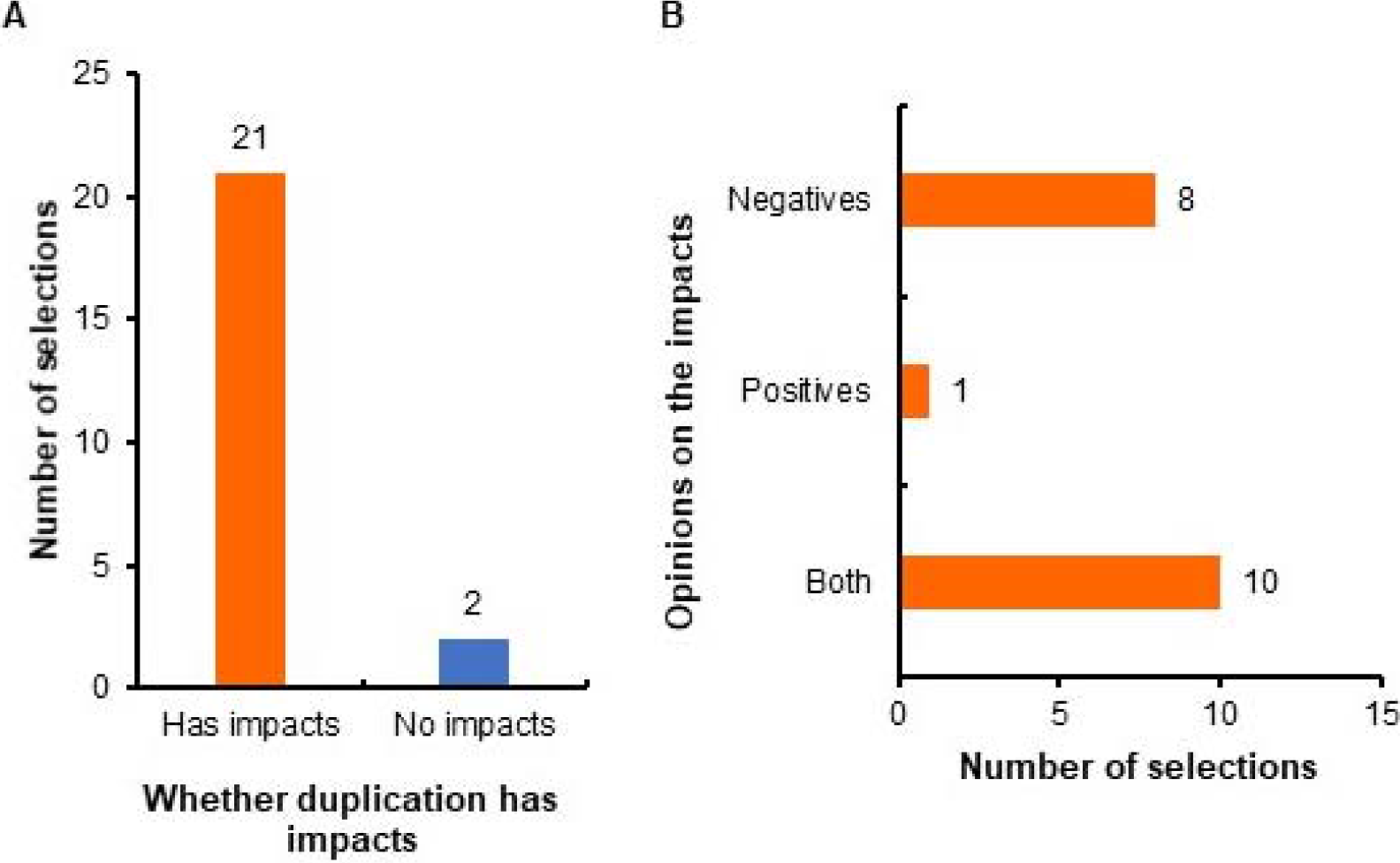
Impacts of duplicate records. A. Do duplicates have impacts? The number of participants who believed whether duplication has impacts or not is shown. B. a more detailed breakdown by type of impact, for those who believed duplication has impacts, is illustrated.

##### Impact on database storage, search and mapping

Representative comments are (1) “When duplicates (sequence only) are in big proportion they will have an impact on sequence search tool like BLAST, when pre-computing the database to search against. Then it’ll affect the statistics on the E-value returned.” [R10], (2) “Duplicates in one resource make exact mappings between 2 resources difficult.” [R21], “Highly redundant records can result in: Increasing bias in statistical analyses; Repetitive hits in BLAST searches.” [R24], and (3) “Querying datasets with duplicate records impacts the diversity of hits and increase overall noise; we have discussed this in our paper on hallmark signatures” [56]. [R8]

##### Impact on meta-analysis in biological studies

Representative comments are (1) “Duplicate transcriptome records can impact the statistics of meta-analysis.” [R1], (2) “Authors often state a fact is correct because it has been observed in multiple resources. If the resources are re-using, or recycling the same piece of information, this statement (or statistical measure), is incorrect.” [R20] (Note that it has been previously observed that cascading errors may arise due to this type of propagation of information [73].) and (3) “Duplicates affect enrichments if duplicate records used in background sets.” [R21]

##### Impact on time and resources

Representative comments are (1) “Archiving and storing duplicated data may just be a waste of resources.” [R12], (2) “Result in time wasted by the researcher.” [R19], and (3) “As a professional curation service; our company suffers from the effects of data duplication daily. Unfortunately there is no pre-screening of data done by Biological DBs and thus it is up to us to create methods to identify data duplication before we commit time to curate samples. Unfortunately, with the onset of next generation data, it has become hard to detect duplicate data where the submitter has intentionally re-arranged the reads without already committing substantial computational resources in advance”. [R9]

Impact on users. Representative comments are (1) “Duplicate records can result in confusion by the novice user. If the duplication is of the ‘low similarity’ type, information may be misleading.” [R19], “Duplicate gene records may be misinterpreted as species paralogs.” [R21], (2) “When training students, they can get very confused when a protein in a database has multiple entries -which one should they use, for example. Then I would need to compare the different entries and select one for them to use. It would be better if the information in the duplicate entries was combined into one correct and more complete entry.” [R23], and (3) “Near identical duplicate records: two or more records are not strictly identical but very similar and can be considered duplicates; because users don’t realise they are the same thing or don’t understand the difference between them.” [R25].

In contrast, practitioners also pointed out two primary positive impacts: (1) identified duplicates enrich the information about an entity; for example, “When you try to look sequence homology across species, it is good to keep duplicates as it allows to build orthologous trees.” [R10] and “When they are isoforms of each other – so while they are for the same entity, they have distinct biological significance.” [R25], and (2) identified duplicates verify the correctness as replications; for example, “On the other hand, if you have many instances of the same data, or near identical data, one could feel more confident on that data point.” [R12] (Note that confidence information ontology can be used to capture “confidence statement from multiple evidence lines of same type” [74].), and “If it is a duplicate record that has arisen from different types of evidence, this could strengthen the claim.” [R13]

The cases outlined above detail the impact of duplication.. Clearly duplication does matter. The negative impacts are broad. They range from databases to studies, from research to training, and from curators to students. The potential impacts are severe: valuable search results may be missed, statistical results may be biased, and study interpretations may be misled. Management of duplication during is a significant amount of labour.

Our survey respondents identified duplicates as having two main positive impacts: enriching the information and verifying the correctness. This has an implicit yet important prerequisite: **the duplicates need to be detected and labelled beforehand**. For instance, in order to achieve information richness, duplicate records must first be accurately identified and cross-references should be explicitly made. Similarly, for confirmation of results, the duplicate records need to be labelled beforehand. Researchers then can seek labelled duplicates to find additional interesting observations made by other researchers on the same entities, that is, to find out whether their records are consistent with others.

The views on *how to manage duplicates* are summarised in **Figure 4**. None of the practitioners regards duplicate detection as unnecessary; 10 practitioners further believe that current duplicate detection methods are not sufficient. We propose the following suggestions accordingly.

**Figure 4.**
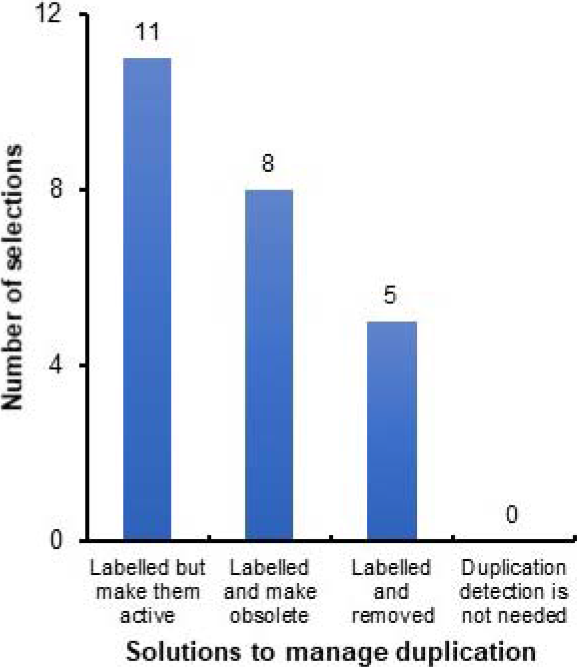
Solutions to duplicate records. How to address duplication? The X-axis represents the options to address duplication; the Y-axis represents the corresponding number of participants selected that option.

##### Precision matters

The methods need to find duplicates accurately: “It should correctly remove duplicate records, while leaving legitimate similar entries in the database.” [R15] and “Duplicate detection method need to be invariant to small changes (at the file level, or biological sample level); otherwise we would miss the vast majority of these.” [R9]

##### Automation matters

In some fields few duplicate detection methods exist: “We re-use GEO public data sets, to our knowledge there is no systematic duplicate detection.” [R7], “Not aware of any software.” [R3] and “I do not use any duplicate detection methods, they are often difficult to spot are usually based on a knowledge of the known size of the gene set.” [R21]

##### Characterisation matters

The methods should analyse the characteristics of duplicates: “A measure of how redundant the database records are would be useful.” [R24]

##### Robustness and generalisation matter

“All formats of data need to be handled cross-wise; it does not help trying to find duplicates only within a single file format for a technology.” [R9]

To our knowledge, there is no universal approach to managing duplication. Similar databases may use different de-duplication techniques. For instance, as sequencing databases, ENCODE uses standardized metadata organisation, multiple validation identifiers, and its own merging mechanism for the detection and management of duplicate sequencing reads; the Sequence Read Archive (SRA) uses hash functions whereas GEO uses manual curation in addition to hash functions [27]. Likewise, different databases may choose different parameters even when using the same de-duplication approach. For instance, protein databases often use clustering methods to handle redundant records. However, the values of chosen similarity thresholds for clustering range from 30% to 100% in different databases [75]. Thus, it is impossible to provide a uniform solution to handling of duplication (as well as other quality issues). We introduce sample solutions used in UniProtKB/Swiss-Prot that demonstrate how quality issues are handled in a single database. The approaches or software used in the UniProtKB/Swiss-Prot curation pipeline may also provide insights into others.

### Beyond duplication: other data quality issues

We also extend the investigation to general quality issues other than duplication, to complement the key insights. We asked the respondents for their opinions on general data quality issues. The two primary questions asked were: *what* data quality issues have been observed in biological databases? and *why* care about data quality? The style is the same as the above questions on duplication. The detailed results are summarized in Supplementary Material File S2. Overall it shows the quality issues can be widespread; for example, each data quality issue has been observed by at least 80% of the respondents.

### Limitations

It is worth noting that while we have carefully phrased the questions in the survey, it may still be the case that different respondents may have different internal definitions of duplicates in mind when responding. For example, some respondents may only consider records with minor differences as redundant records whereas others may also include records with larger differences, even though they selected the same option. We acknowledge that this diversity of interpretation is inevitable – data is multifaceted; hence so is data quality and the associated perspectives on it. The internal definitions of duplicate records depend on more specific context and there is indeed no universal agreement [24]. However, we argue that this does not detract from the results of the survey; respondents provided clear examples to support their choices and those examples demonstrate that the duplicate types do impact biological studies, regardless of internal variation in specific definitions. Such internal differences are also observed in other data quality studies, such as reviews on general data quality [76] and detection of duplicate videos [77].

It is also noteworthy that some databases primarily serve an archival purpose, such as INSDC and GEO. The records in these databases are directly coordinated by record submitters; therefore, the databases have had relatively little curation compared to databases like UniProtKB/Swiss-Prot. Arguably, data quality issues are not major concerns from an archival perspective. We do not examine the quality issues in archival databases; rather, we suggest labelling duplicate records or records with other quality issues (without withdrawing or removing the records) could potentially facilitate database usage. The archival purpose does not limit other uses; for example, studies including BLAST searches against GenBank for sequence characterization [78–80]. In such cases, the sequences and annotations would impact the related analyses.

However, quality issues may be important in archival databases. Indeed, in some instances the database managers have been aware of data quality issues and are working on solutions. A recent work proposed by the ENCODE database team concerns the quality issues, in particular duplication in sequencing repositories such as ENCODE, GEO and SRA [27]. They acknowledge that, while archival databases are responsible for data preservation, duplication affects data storage and could mislead users. As a result, they propose three guidelines to prevent duplication in ENCODE and summarise other de-duplication approaches in GEO and SRA; furthermore, the ENCODE work encourages making a community effort (such as archival databases, publishers, and submitters) to handle quality issues.

### Biocuration: a solution to data quality issues in biological databases

In this section, we introduce solutions to data quality issues in biological databases. Biocuration is a general term that refers to addressing data quality issues in biological databases. We provide a concrete case study on the UniProtKB/Swiss-Prot curation pipeline – consisting of a detailed description on the curation procedure and an interview with the curation team leader. It provides an example of a solution to different quality issues.

### The curation pipeline of UniProtKB/Swiss-Prot

UniProtKB has two data sections: UniProtKB/Swiss-Prot and UniProtKB/TrEMBL. Sequence records are first deposited in UniProtKB/TrEMBL and then selected records are transferred into UniProt/Swiss-Prot. Curation in UniProtKB has two stages: (1) automatic curation in UniProt/TrEMBL, where records are curated by software automatically without manual review, and (2) expert (or manual) curation in UniProtKB/Swiss-Prot on selected records from UniProtKB/TrEMBL. A major task in automatic curation is to annotate records using annotation systems; for example, UniRules, which contains rules created by biocurators, and external rules from other annotation systems, such as RuleBase [81] and HAMAP [82], are used in this task. Rule UR000031345 is an example of UniRules (http://www.uniprot.org/unirule/UR000031345); Record B1YYB is also a sequence record example that was annotated using the rules during automatic curation. For expert curation, biocurators run a comprehensive set of software, search supporting information from range of databases, manually review the results and interpret the evidence level [31]. **Table 1** describes representative software and databases used in expert curation [14, 83–98]. This expert curation in UniProtKB/Swiss-Prot has 6 dedicated steps, shown in Table 1 and explained below.

**Table 1.** Representative software and resources used in expert curation. *Note*: A complete set of the software, including the detailed versions of the software, can be found in UniProt manual curation standard operating procedure documentation (www.uniprot.org/docs/sop_manual_curation.pdf).

| Curation steps | Software/<br>Databases | Purpose | Ref. |
| --- | --- | --- | --- |
| **Sequence curation** |  |  |  |
| Identify homologs | BLAST | Sequence alignment | [83] |
|  | Ensembl | Phylogenetic resources | [84] |
| Document inconsistencies | T-Coffee | Analysis of causes of | [85] |
|  | Muscle | inconsistencies due to | [86] |
|  | ClustalW | duplication | [87] |
| **Sequence analysis** |  |  |  |
| Predict topology | Signal P | Signal peptides prediction | [88] |
|  | TMHMM | Transmembrane domain prediction | [89] |
| Post-translations | NetNGlyc | N-glycosylation sites prediction | [90] |
|  | Sulfinator | Tyrosine sulfation sites prediction | [91] |
| Identify domains | InterPro | Retrievals of motif matches | [92] |
|  | REPEAT | Identification of repeats | [93] |
| **Literature curation** |  |  |  |
| Identify relevant literature | PubMed | Literature resources | [94] |
|  | iHOP |  | [95] |
| Text mining | PTM | Information extraction | [96] |
|  | PubTator |  | [97] |
| Assign GOs | GO | Gene ontology terms | [14] |
| **Family curation** | Same as identify homologs |  |  |
| **Evidence attribution** | ECO | Evidence code ontology | [98] |

#### Sequence curation

This step focuses on de-duplication. It has two components: (1) Detect and merge duplicate records. (2) Analyse and document the inconsistencies caused by duplication. In this specific case ‘duplicates’ are records belonging to the same genes: an example of entity duplicates. Biocurators perform BLAST searches and also search other database resources to confirm whether two records are the same genes, and merge them if they are. The merged records are explicitly documented in the record’s *Cross-reference* section. Sometimes the merged records do not have the same sequences, mostly due to errors. Biocurators have to analyse the causes of those differences and document the errors.

#### Sequence analysis

Biocurators analyse sequence features after addressing duplication and inconsistencies. They run standard prediction tools, review and interpret the results, and annotate the records. The complete annotations for sequence features cover 39 annotation fields under 7 categories: Molecule processing, Regions, Sites, Amino acid modifications, Natural variations, Experimental info, and Secondary structure (http://www.uniprot.org/help/sequence_annotation).

As such, it involves a comprehensive range of software and databases to facilitate sequence analysis, some of which are shown in Table 1.

#### Literature curation

This step often contains two processes: retrieval of relevant literature and application of text mining tools to analysis of text data, such as recognising named entities [99] and identifying critical entity relationships [100]. The annotations are made using controlled vocabularies (the complete list is in the UniProt keyword documentation via http://www.uniprot.org/docs/keywlist) and are explicitly labelled as “*Manual assertion based on experiment in literature*”. Record Q24145 is an example that was annotated based on findings published in literature (http://www.uniprot.org/uniprot/Q24145).

#### Family-based curation

This step transitions curation from single-record level to family-level, finding relationships amongst records. Biocurators identify putative homologs using BLAST search results and phylogenetic resources and make annotations accordingly. The tools and databases are the same as those in the *Sequence curation* step.

#### Evidence Attribution

This step standardises the curations made in the previous steps. Curations are made manually or automatically from different types of sources, such as sequence similarity, animal model results and clinical study results. This step uses the Evidence and Conclusion Ontology (ECO) to describe evidence in a precise manner: it details the type of evidence and the assertion method (manual or automatic) used to support a curated statement [98]. As such, database users can know how the decision was made and on what basis. For example, ECO_0000269 was used in the literature curation for Record Q24145.

##### Quality assurance, integration and update

The curation is complete at this point. This step finally checks everything and integrates curated records to the existing UniProtKB/Swiss-Prot knowledgebase. Those records will then be available in the new release. In turn, it helps further automatic curation within UniProtKB/Swiss-Prot. The newly made annotations will be used as the basis for creating automatic annotation rules.

### The curation in UniProtKB/Swiss-Prot: an interview

We interviewed UniProtKB/Swiss-Prot annotation team leader Sylvain Poux. The interview questions covered how UniProtKB/Swiss-Prot handles general data quality issues. Some of the responses are also related to specific curation process in UniProtKB/Swiss-Prot which shows that the solutions are database-dependent as well. The detailed interview is summarized in the Supplementary Material File S3. We have edited the questions for clarity, and omitted answers where Poux did not offer a view.

The above case study demonstrates that biocuration is an effective solution to diverse quality issues. Indeed, since 2003, when the first regular meeting amongst biocurators was held [101], the importance of biocuration activities has widely been recognised [20, 102–104]. Yet, on the other hand, the biocuration community still lacks broader support. A survey of 257 former or current biocurators showed that biocurators suffered from a lack of secured funding for primary biological databases, exponential data growth, and underestimation of the importance of biocuration [69]; consistent results were also demonstrated in other studies [105, 106]. According to recent reports, the funding for model-organism databases will be cut 30%–40% and the same threat applies to other databases [107–109].

## Conclusion

In this study, we explored the perspectives of both database managers and database users on the issue of data duplication – one of several significant data quality issues. We also extended the investigation to other data quality issues to complement this primary focus. Our survey of individual practitioners showed that duplication in biological databases is of concern: its characteristics are diverse and complex, its impacts cover almost all stages of database creation and analysis, and methods for managing the problem of duplication, either manual or automatic, have significant limitations. The overall impacts of duplication are broadly negative, and the positive impacts such as enriched entity information and validation of correctness rely on the duplicate records being correctly labelled or cross-referenced. This suggests a need for further development of methods for precisely classifying duplicate records (accuracy), detecting different duplicate types (characterisation), and achieving scalable performance in different data collections (generalisation). In some specific domains duplicate detection software (automation) is a critical need.

The responses relating to general data quality further show that data quality issues go well beyond duplication. As can be inferred from the survey we conducted, curation – dedicated efforts to ensure that biological databases represent accurate and up-to-date scientific knowledge – is an effective tool for addressing quality issues. We provide a concrete case study on the UniProtKB/Swiss-Prot curation pipeline as a sample solution to quality issues. However, manual curation alone is not sufficient to resolve all data quality problems due to rapidly growing data volumes in a context of limited resources. A broader community effort is required to manage data quality and to provide support to facilitate data quality and curation.

## Authors’ contributions

QC, JZ and KV initiated the survey, analysed the results and wrote the paper. RB, IE, CJ, AL, MM, JO, MR, JS, and RY contributed to presenting the views and revising the paper. All authors read and approved the final manuscript.

## Acknowledgments

The project receives funding from the Australian Research Council through a Discovery Project grant, DP150101550. We thank Sylvain Poux for contributions to the UniProtKB/Swiss-Prot curation case study. We acknowledge the participation of the following people in the survey: Cecilia Arighi (University of Delaware), Ruth C Lovering (University College London), Peter McQuilton (University of Oxford), and Valerie Wood (University of Cambridge).

## Supplementary material

**File S1 Survey questions**

**File S2 Results and discussions on quality issues beyond duplication**

**File S3 UniProtKB/Swiss-Prot annotation team leader interview details**

